## Supplementary Information for "*improv*: A software platform for real-time and adaptive neuroscience experiments"

### *improv* installation

**System requirements.** *improv* was designed to be as system-agnostic as possible. The core functionality is written for Python 3.10, but is compatible for Python 3.7 – 3.10, and requires the installation of a few other packages (such as numpy, pyarrow, and pyzmq). Installation of *improv* has been tested and confirmed on Linux (Ubuntu), Mac OS X, and Windows running Windows Subsystem for Linux (WSL).

**Base installation.** We provide complete installation instructions in the Readme file on the Github repository and through our documentation website: https://project-improv.github.io/improv/installation.html.

The easiest method for installation is through the Python Package Index PyPI:

```
pip install improv
```

Users can also use a local clone from Github to install the package, especially if they desire to edit or contribute to the codebase. Briefly, two calls are needed:

1. `python -m build` to build the package with all requirements,
2. `pip install -e .` to install the package inside the local directory.

After either installation method, *improv* can be executed on a particular user-configured setup with:

```
improv run experiment.yml
```

### *improv* software design

Configuration in *improv* is streamlined and simple, requiring only that a user define (1) what processing steps (i.e., actors) are part of the pipeline, and (2) in what order they should be executed. The pipeline definition is lightweight: directed graphs are specified by text-based configuration files with two sections (**Extended Data Fig. 1**). In the first section, actors are specified by the names of their respective Python classes, with any additional information passed as keyword arguments to class constructors. In the second section, connections among actors are specified by listing the consumers for each actor's outputs. On startup, *improv* handles construction for each of these queues and links them with their respective actors. *improv* can handle both single and multiple output queues to accommodate arbitrary configurations.

Importantly, all instantiation, memory-sharing, execution flow, and logging are handled by *improv*, leaving the user free to focus on code within an actor. Furthermore, any Python code (or code executable in a Python environment) is acceptable within an actor: we do not supply a library of functions or require that a user follow particular implementation. Thus, almost anything a user wants to compute or run within an actor is available, including integration with other software tools.

**Extended Data Fig. 1b** illustrates this for a simple Processor actor. Here, users define the runStep method which is run in a rapid and infinite loop by *improv*. The method executes code to receive a data store ID (key) from the input queue and uses it to retrieve the datum (in this example, an array called self.frame of neural activity). It computes the frame average and places the result back into the data store. The keys to identify those results are then published in the supplied output queue self.q_out. In our example pipeline, these are retrieved for display by the visual actor (the graphical user interface). This actor can remain unaware of all other aspects of the experiment and awaits only the next piece of data to execute its computation.

*improv* only specifies the semantics of data pipelines, freeing users from the need to manage implementation details. This is facilitated by two key components: a shared, in-memory data store, discussed above, and a central controller program, dubbed Nexus. Once users have defined each processing step in the pipeline and the dependency relationships between them, Nexus is responsible for the actual orchestration of experiments. At runtime, Nexus instantiates each actor, configures information and execution flow, and handles both error signaling and user interactions. On startup, it takes in a list of desired actors and associated attributes from the configuration file, instantiates each class, and provides it access to the shared data store. *improv*'s user contract is thus both extensible and simple: users write custom Python classes for any new analyses they require, but they do not need to know the internals of classes implementing other pipeline steps.

Once the experimental pipeline is started, Nexus executes each class in a separate process and monitors its progress. Each process corresponding to an actor is kept alive continuously, waking as data become available in its input queue. In this way, the system leverages concurrency to overcome delays due to serial processing or input/output overhead. Communication is handled asynchronously, using a custom class combining Python's asyncio and multiprocessing libraries. As a result, the entire pipeline is robust to failures: no one actor or internal task can suspend the system, which would effectively cause the experiment to terminate. In addition, each object placed into the data store and every parameter change can be logged to disk, effectively creating a snapshot of the system state at each moment in the experiment. This ensures a robust audit trail, such that any data or associated analysis can be reproduced by later offline analyses.

### API and Sample use cases

**Graphical user interface.** For our specific experimental integration of *improv* with a two-photon calcium imaging setup, we also constructed a simple graphical user interface (GUI) to provide user control and real-time updates of all images and analyses (**Supplementary Video 1**). For the paradigm described in the main text, we plot fluorescence or extracted spikes as a function of the last 500 frames (scrolling window) for both the population average and a user-selected neuron. Response profiles are plotting as circular tuning curves adjacent to those line plots. Raw images acquired from the setup are shown to the left, and the processed and analyzed frame is shown to the right. Neurons in the processed frame are colored by their tuned responses and can be selected via mouse click by a user to display its data above. On the right side we display the online results of the LNP model fit. The top plot shows the negative log-likelihood function being minimized as more frames are analyzed, and the bottom plot displays the inferred weight matrix of connections among the top 10 neurons with largest model weights. Again, by selecting a neuron in the center processed frame plot, the connections associated with that neuron are displayed as green lines.

**Integration with other tools.** While *improv* represents the only fully extensible tool dedicated in the online setting, there are a host of excellent tools available for offline analysis. As a step toward integration with these tools, we have implemented proof-of-concept interfaces between *improv* and both Suite2p (via its Python command-line interface) and ScanBox (via the MATLAB execution environment) (**Extended Data Fig. 5**). Other pipeline or workflow generating software packages have also tackled this problem of efficient pipelining, even specifically for the neuroscience community (Gorgolewski, 2011). Our system by contrast does not attempt to directly bundle collections of applications for a multi-use toolbox, but instead is a lightweight scaffolding approach that provides containers (actors) to accommodate most any application the user needs. This also ensures our code does not fall into maintenance traps, but rather easily enables updates or adding new functionality.

We additionally looked at integration across different programming languages. For instance, both Python and Julia have packages designed for interfacing with one another. Using the Python package PyJulia, we can compile and execute a piece of Julia code from within a Python program. Combining this with the Julia package PyCall to do the reverse, we can flexibly transfer data from *improv* to Julia for quick analysis (e.g., gradient descent of an LNP model) and transfer the results back into *improv* once again. Notably, this transfer between languages can be accomplished with time-efficient no-copy wrappers if using NumPy arrays. For an example implementation using Julia see the julia branch at github.com/pearsonlab/improv.

### Benchmarking

Our optimized system proved capable of preprocessing, analyzing, and visualizing results at rates faster than a simulated acquisition frequency of 30 Hz, which is roughly 6.7 MB/s for one test data set. Moreover, we show that even faster rates than this are possible. Whereas a serial implementation of the analysis exceeds the per cycle time budget of 33ms, *improv*'s inbuilt computational concurrency accomplishes all the same steps well within these time constraints, suggesting that this approach can also scale to volumetric light-sheet microscopy, e.g., for the entire zebrafish brain at 0.8 Hz for 41 planes (roughly 7 MB/s) (Ahrens, 2013). And while short periods of heightened preprocessing time can occur, any lag between acquisition and final analysis quickly dissipates. In fact, *improv* was able to process data continuously for more than twenty-four hours without crashing or accumulating more than a single frame of processing lag (**Extended Data Fig. 2**). Furthermore, concurrency allows *improv* to easily ignore "bad actors," i.e., missing or slow frames that frequently lag or crash, and maintain the processing the pipeline. To illustrate system stability, we simulated an imaging experiment in which the 'acquisition' actor drops frames: rather than await missing frames that may never appear (suspending the experiment) or raising errors (halting the experiment), each actor continues its own execution loop, processing any future data when they become available. Importantly, once acquisition resumes, the entire system recovers automatically, without delays or increased processing time.

### Supplementary Videos

**Supplementary Video 1 |** A screen recording of our GUI used in **Fig. 2** to conduct our simulated experiments. Similar GUIs were used for *in vivo* experiments conducted for **Fig. 5** and **Fig. 6**.

**Supplementary Video 2 |** Online regression coefficients overlaid back onto the original behavioral video of the mouse's face and paws for the experiment described in **Fig. 3**. Color intensity corresponds to instantaneous magnitude of the coefficients across the image.

**Supplementary Video 3 |** Movie generated from data acquired live during a Bayesian optimization experiment described in **Fig. 5**. (Left) Estimated calcium fluorescence traces for each stimulus window. (Center, Right) The Gaussian process tuning curve and associated uncertainty, respectively, for each stimulus displayed. The overlaid dots denote the sampled location (which visual stimulus) with color corresponding to the empirical neural activity in response to that stimulus.

**Supplementary Video 4 |** Example two-photo calcium fluorescence recording during visual stimulation and photostimulation for the experiments conducted in **Fig. 6**. The top left overlay shows the visual stimuli displayed or the time of photostimulation with a red dot. An increase in neural activity signal is seen directly following visual stimuli or photostimulation events.

**Supplementary Video 5 |** A screen recording demonstrating the text interface supplied with *improv* and basic usage. An experiment is loaded from a .yaml file, initialized with the setup command, then run, stopped, and quit using typed commands. Log messages are displayed in the top portion of the window, with user input accepted in the below portion.
